# Early life exploration behavior and life-history loci are co-localized in an adaptive genomic hotspot in Atlantic salmon

**DOI:** 10.1101/2025.05.28.656530

**Authors:** Tutku Aykanat, Jaakko Erkinaro

**Affiliations:** Organismal and Evolutionary Biology Research Program, Faculty of Biological and Environmental Sciences, University of Helsinki, Finland; Natural Resources Institute Finland (Luke), Oulu, Finland

**Keywords:** Correlated traits, exploration, habitat shift, age at maturity, epistasis, pleiotropy, Atlantic salmon

## Abstract

1. The traits that are important for adaptation may exhibit genetic correlation due to pleiotropy or as a result of linkage. Understanding the genetic architecture of such correlations is important for predicting the selection response of populations. Exploration in fishes is a behavioral trait by which individuals may find habitats with better foraging and growth opportunities that subsequently improve their fitness. For example, in Atlantic salmon (*Salmo salar)*, all individuals originate in spawning rivers where females lay eggs, but some juveniles migrate to small tributaries which are not spawning areas but provide favorable habitat patches for young salmon. The increased growth in these nursery streams may facilitate earlier sexual maturation, implying a potential correlation between exploration and maturation traits.
2. In this study, by sampling juveniles from two wild populations in the large Teno River catchment in northernmost Fennoscandia, we tested the genetic association between exploration behavior with nursery streams across four SNPs that span a 70-kb long genomic region with a major effect on age at maturity variation. Three of these SNPs are missense mutations in the *vgll3* and *akap11* genes, and one SNP tags a putative regulatory region with the strongest association with the age-at-maturity trait.
3. We show that the exploration behavior was linked to the genomic region in one of the two studied populations. However, the genetic association was substantial in the missense SNP located in the *akap11* gene, which is farthest away from the *vgll3* SNPs and previously ruled out as being linked to the age at maturity. We also detected a marginal interaction effect between SNPs in the *vgll3* gene and *akap11*, indicating a potentially complex genetic architecture underlying the trait variation.
4. Our results suggest that exploration and age at maturity are co-inherited within the same haplotype block, but we find no evidence for direct causality. These two traits may form a co-adapted trait complex that may be instrumental in local adaptation processes.

## Introduction

Elucidating the genetic architecture of adaptive traits is essential to predict the evolutionary trajectories linked to natural selection in the wild. While genome-wide scans are essential for locating genomic regions associated with phenotypic variation or adaptation, their utility to assess fine-scale adaptive landscape is limited, especially in natural populations. For example, a genomic region may harbor multiple loci associated with trait variation and, depending on the number of loci and the linkage disequilibrium between them, the predicted response to selection will vary. Close genomic proximity or close genetic linkage distances between adaptive genetic loci can constrain adaptation (Connallon & Hall 2018), but may also be selected for to preserve co-adapted allele combinations in the form of epistatic interaction (Yeaman 2013; Bank 2022). For example, the evolution of inversions reduces recombination along the inversion and allows gene complexes to be preserved over large genomic distances (Kirkpatrick 2010; Küpper et al. 2015). Such an epistasis has been suggested as an important form of genetic architecture that facilitates the coevolution of coadapted trait complexes, such as behavior and reproduction or behavior and life history (Küpper et al. 2015; Tuttle et al. 2016; Funk et al. 2021).

Likewise, population models suggest such nonrecombining regions may be preserved even without a need for coadaptation or epistasis (Kirkpatrick & Barton 2006; Hoffmann & Rieseberg 2008), suggesting their potential prevalence in the natural populations.

Loci that are in close proximity have inherently low recombination rate. Hence, similar to long nonrecombining genomic regions, they could also harbor coadapted gene complexes or independent adaptive loci. However, the prevalence of such mechanisms is not well understood in natural populations. For example, it is challenging to distinguish whether multiple traits are genetically associated with independent loci within the region (spurious pleiotropy) or due to true pleiotropy (linked to one causal genic polymorphism, see, e.g., Solovieff et al. 2013). Distinguishing these two modes of inheritance has implications. In the evolutionary sense, pleiotropic loci are harder to decouple than tightly linked independent loci (Chebib & Guillaume 2021) hence may be less responsive to evolutionary pressures. On the other hand, synergistic pleiotropy may evolve as a result of strong natural selection, in which case mutations effecting multiple fitness traits may be selected for (McGee et al.2016). Likewise, multivariate phenotypes associated with tightly linked loci might indicate coadaptation (between traits) within local demes (Yeaman & Whitlock 2011), while phenotypes under a pleiotropic effect of a gene may have a developmental basis (Wagner, Pavlicev & Cheverud 2007). Therefore, linking genetic architecture to ecological processes and adaptation requires information not only on the phenotypes associated with the genomic region, but also on the fine-scale genetic association between phenotypes and polymorphisms within genomic regions.

The *vgll3* locus in Atlantic salmon (*Salmo salar*) has strong potential as a genomic tool to better understand the adaptive dynamics in the wild associated with a small linked genomic region. The *vgll3* genomic region, which spans approximately 70 kb and encompasses the *vgll3* and *akap11* genes, was initially identified as a major genomic region that explains age at maturity, a life history trait with substantial fitness effects (Ayllon et al. 2015; Barson et al. 2015). Further research identified that variation in other traits was also linked with genetic variation in the region, such as precocious male maturation (Debes et al. 2021), repeat spawning (Aykanat et al. 2019), metabolism (Prokkola et al. 2022; Prokkola et al. 2024), foraging behavior (Aykanat et al. 2024), exploration and aggression (Bangura et al. 2022; Bangura et al. 2024), morphology (Aykanat et al. 2025) and early life fitness (Aykanat et al. 2024). The region harbors a large number of single nucleotide polymorphisms (SNP), as revealed by whole genome sequencing (e.g., Barson et al. 2015), but four SNPs remain the most important candidates for the association of phenotypes. These include two potentially significant functional missense polymorphisms within the coding regions in the *vgll3* gene (*vgll3*Met54Thr and *vgll3*_Asn323Lys_), one missense polymorphism in the *akap11* gene (*akap11*_Val214Met_), and a putatively regulatory SNP with the strongest association with maturation age, so-called *vgll3*_TOP_ (Barson et al. 2015). However, in most of these studies, the trait-gene association was not further refined. For example, in studies that involved cross-breeding designs, parental fish were usually selected to have a homogeneous *vgll3* haplotype block with full linkage among these four important SNPs, primarily to increase the inference power to detect the additive genetic relation. In one exceptional case, cross-breeding was designed to segregate the genetic variation between the *vgll3* and *akap11* genes, which allowed further refinement of the location of causal SNPs within the genomic region (Sinclair-Waters et al. 2022). Genetic variation across tightly linked loci can also be refined in wild populations by exploiting natural segregation within the region. For example, Aykanat et al. (2019) attempted to narrow down the genetic variation in the *vgll3* region associated with the repeat spawning phenotype along four important SNPs in the region, but none of the four SNPs explained the phenotypic variation better than the others, indicating insufficiency of power. Taken together, the presence of multiple adaptive traits and putatively functionally important SNPs linked to the genomic region signify the importance of better characterizing the genetic architectures. This will provide insight into the dynamics of evolutionary change of the genomic region within and among Atlantic salmon populations.

Exploration in fishes is an important behavioral trait, enabling populations to spread and occupy new habitat patches and resources. It can reduce the negative effects of density dependence (Einum, Sundt‐Hansen & H. Nislow 2006), and improve population stability (Baker et al. 2025). In juvenile Atlantic salmon, exploration within freshwater (before emigration to the ocean) has the potential to contribute to improved feeding opportunities, better growth, and ultimately higher survival (Erkinaro & Niemelä 1995; Erkinaro & Erkinaro 1998; Johansen, Elliott & Klemetsen 2005a; Johansen, Elliott & Klemetsen 2005b). Such habitat shifts may also affect individual life history patterns (Erkinaro et al. 1997) via improved feeding and growth opportunities (e.g., Udea et al. 2025), making the individuals more likely to exceed the maturation threshold (Metcalfe 1998; Thorpe et al. 1998; Jonsson & Jonsson 2014). At the genetic level, exploration and covarying traits such as metabolism and aggression have been linked with the major loci affecting age at maturity, *vgll3* (Bangura et al. 2022; Prokkola et al. 2022; Bangura et al. 2024; Prokkola et al. 2024). For example, using a common garden design in an aquarium setting, Bangura et al. (2024) showed that juvenile Atlantic salmon with the late-maturing *vgll3*^LL^ genotype (L referring to the allele associated with late maturation) were significantly more active in the 30-minute period after their release into a novel environment than individuals with the early-maturing *vgll3*^EE^ genotype (E referring to the allele associated with early maturation). This indicates that exploration and maturation could be influenced by the same genetic locus. However, in the cross-design, the *vgll3* haplotype block was in complete linkage, and the study could not further refine the genetic basis within the genomic region.

Traditionally, Atlantic salmon juveniles are considered relatively site-attached and sedentary in their early years in freshwater (see, e.g., Jonsson & Jonsson 2011). However, in some river systems, older age groups of juvenile salmon (parr) disperse to more favorable habitat patches within freshwater systems, including lacustrine areas and smaller tributaries (Erkinaro 1995; Erkinaro, Julkunen & Niemelä 1998; Halvorsen & Jørgensen 2006).

These small tributaries are nursery streams that are not suitable for salmon reproduction but may provide favorable growth and survival conditions for salmon parr by means of increased food availability and decreased competition and predation (Erkinaro & Niemelä 1995; Erkinaro & Erkinaro 1998; Erkinaro, Julkunen & Niemelä 1998; Johansen, Elliott & Klemetsen 2005b). The extent of exploratory behavior to nursery streams and its adaptive importance may be influenced by various factors, including environmental conditions, genetic predispositions, and population density (e.g., Erkinaro et al. 1997; Johansen, Erkinaro & Amundsen 2010). For example, the use of nursery streams by juvenile Atlantic salmon has been observed especially in northern areas with depauperate fish fauna that exhibit less competition and predation threat to juvenile salmon (Erkinaro 1995; Erkinaro & Gibson 1997; Shustov, Baryshev & Belyakova 2012).

Measuring the active migration of juvenile Atlantic salmon to nursery streams is a good proxy for quantifying exploration behavior in natural settings (Erkinaro, Julkunen & Niemelä 1998) and may help us to quantify genetic basis of the exploration across populations. Furthermore, such migratory behavior may be linked to maturation age during adulthood. The relative proportion of early maturation is shown to be higher among individuals that occupy nursery streams during the juvenile stage (Erkinaro et al. 1997), which may be facilitated by improved growth opportunities and the likelihood of reaching the maturation threshold (e.g., Thorpe et al. 1998). Thus, a genetic covariation between exploration and maturation timing may be hypothesized as the basis of this observation.

Here, we quantified the genetic basis of exploration to nursery streams in two wild Atlantic salmon populations in the Teno River system in northernmost Fennoscandia. We first ask whether the *vgll3* genomic region is associated with exploratory behaviors to nursery streams. By further exploring the genetic architecture within the *vgll3* haploblock, we asked if exploration and age at maturity traits are pleiotropic or associated with different genetic loci, and if epistasis within the region may play a role in shaping trait variation.

## Material and Methods

### Study area and sampling

The Teno River (Deatnu in Sámi, Tana in Norwegian) catchment is located in far-north Europe (68–70°N, 25– 27°E), runs northward between Finland and Norway, and drains into the Barents Sea (Supplementary Figure 1). The Teno system supports a number of Atlantic salmon populations with wide diversity in life history characteristics and genetic structure within and among populations (Vähä et al. 2017; Erkinaro et al. 2019). Juvenile salmon samples used in this study were originally collected for a series of studies characterizing the exploration behaviors of juvenile salmon from the main stem spawning areas into small nursery streams in relation to environmental conditions and fish age (Erkinaro & Niemelä 1995; Erkinaro, Julkunen & Niemelä 1998). Samples were collected from two populations that reproduce in the Teno main stem and Inarijoki (Anárjohka in Sámi and Norwegian), a large headwater tributary of the Teno River. These two populations exhibit a low level of genetic divergence (*F*_ST_ = 0.012, Vähä et al. 2008; Aykanat et al. 2015), but have substantial differences in their life histories, with the former population having high proportions of large, later-maturing individuals, and the latter population primarily consisting of younger individuals at maturity (Czorlich et al. 2018, Erkinaro et al. 2019). In the spawning rivers, the Teno main stem and Inarijoki, where adult Atlantic salmon reproduce and females lay eggs, juvenile salmon were captured using 600-to 900-V pulsed DC electrofishing equipment. Scales were sampled (initially for age determination, but later used for DNA extraction) and after sampling, the fish were released unharmed at the site of capture.

The sampling sites in both spawning rivers were located in close proximity (up to 100 meters) to the outlets of two nursery streams, Baððá and Guoldnájohka (Supplementary Figure 1), which drain into the Teno main stem and Inarijoki, respectively (Erkinaro, Julkunen & Niemelä 1998). Juvenile salmon in the streams were captured with a counting fence installed at the lowest parts of the streams, 20–50 m from the outlets, and as above, the scales were sampled. After sampling, fish were released in the direction in which they had been heading (*cf*., Erkinaro, Julkunen & Niemelä 1998). Migrant fish were classified as smolt or parr based on external characteristics, especially coloration, visibility of parr marks, and size (Erkinaro & Niemelä 1995). Overall, we sampled and genotyped 785 individuals, of which 635 were used in the main analysis after filtering out IDs with missing genotype information in focal SNPs (see below) or missing cohort year information (i.e., due to poor-quality scale samples; Supplementary Table 1).

### DNA extraction, SNP genotyping by targeted sequencing

DNA was extracted from juvenile Atlantic salmon scales using QIAamp 96DNA QIAcube HTKit (Qiagen) following the manufacturer’s protocol. Individuals were genotyped by targeted sequencing using a SNP panel consisting of 173 SNP markers using a GTSeq approach (Campbell, Harmon & Narum 2015) as described in (Aykanat et al. 2020). The SNP panel contained four SNPs from the *vgll3* genomic regions including the SNP that is most highly associated with age at maturity (*vgll3*_TOP_), two missense mutations in the *vgll3* gene (*vgll3*_Met54Thr_ and *vgll3*_Asn323Lys_), and one missense mutation in the *akap11* gene (*akap11*_Val214Met_, Barson et al. 2015). These SNPs are located on chromosome 25 and are 0.8 kb (*vgll3*_Met54Thr_), 2.9 kb (*vgll3*_Asn323Lys_), 10.9 kb (*vgll3*_TOP_), and 65.5 kb (*akap11*_Val214Met_) downstream of the transcription start site of the *vgll3* gene. The SNP panel also included *six6* locus, a SNP highly spatially divergent between Atlantic salmon populations (Barson et al. 2015), and the so-called outlier and baseline SNPs. The outlier module includes 53 highly differentiated SNPs between the Inarijoki and Teno main stem populations, allowing to confirm the population of origin of individuals between these two closely related populations (data not shown). The baseline module included 136 putatively neutral markers in low linkage disequilibrium, distributed throughout the whole genome and previously filtered to have a minor allele frequency greater than 0.05 and heterozygosity greater than 0.2 for these two populations (Aykanat et al. 2016). Individuals who did not have genotype information on any of these SNPs were excluded from the analysis. Bonferroni correction was applied when multiple SNPs were compared.

### Statistical analysis

We first tested if the exploration trait is colocalized in the *vgll3* genomic region by comparing single SNP association to exploration behavior, which may also address whether exploration and age at maturity are pleiotropic or associated with different loci within the region. We modeled this separately for the Teno main stem and Inarijoki populations using a generalized linear mixed model with a binomial error structure as follows:

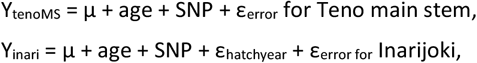

where Y is a Boolean response variable for the explorative behavior of individuals, in which individuals sampled in the main stem were assumed to be sedentary, and coded as “0”, and individuals sampled in the streams, either descending or ascending, were assumed to be explorative and coded as “1”. SNP is the genotype score of SNPs, coded as 0 when individuals are homozygous for the reference alleles, 1 when homozygous for the alternative alleles, and 0.5 when heterozygous. Age of the fish was determined from the sampled scales. For the Inarijoki population, ε_hatchyear_ is the random variance associated with the hatch year (i.e., cohort year) calculated as the year sampled minus the age of the fish. Hatch year controls for potential cohort specific allele frequency differences between different years. Neither the hatch year nor the sampling year overlapped for the Teno main stem populations between nursery streams and main stem collections, so for this population, the model did not include the random term ε_hatchyear_. The model was implemented for four focal SNPs in the vgll3 genomic region (*vglll3*_TOP_, *vgll3*_Met54Thr_, *vgll3*_Asn323Lys_, *akap1*_Val214Met_),, *six6 locus* that is shown to be associated with age at maturity and metabolism (Sinclair-Waters et al. 2020; Prokkola et al. 2022; Prokkola et al. 2024), and the rest of 169 SNPs in the SNP panel to assess baseline association levels in the putatively unrelated loci. Using a similar model structure as above, we then implemented a set of models to quantify the effect of epistasis within the genomic region by modeling pairwise interactions of SNPs within the *vgll3* genomic regions:

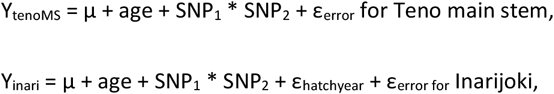

Whereby, SNP_1_ and SNP_2_ are two of the four focal SNPs, and we all possible models that included any pairwise comparisons between them.

Finally, we compared the differences in gene trait association between populations, i.e., Inarijoki and Teno main stem, which are in close proximity and genetically similar, but with substantial differences in their life histories. Such a difference may potentially indicate gene-by-environmental, gene-by-gene (i.e., due to populations genetic background), or a combination of these two effects.

All models were run using the glmmTMB package, version 1.1.5, (Brooks et al. 2017) in R software version 4.4.1 (R Core Team 2019), using binomial error structure. Marginal means for genotype effects and their confidence intervals were calculated using the emmeans package, version 1.8.5 (Lenth, 2023). Model diagnostics were performed using the DHARMa package, version 0.4.6 (Hartig 2022).

## Results

Overall, the genotyping success rate was generally low (85%, Supplementary Figure 2), mainly due to the small amount of material available for DNA extraction. After removing individuals with no genotype success in any of the focal SNPs (N=143) and those with no scale aging data (N=15, of which eight also did not have genotype success), 635 individuals remained in the final dataset out of an initial 785.

In the Teno main stem population, exploration behavior towards the nursery stream, Baððá, was not significantly associated with any of the four SNPs in the *vgll3* genomic region, or any other SNPs in the panel after the Bonferroni correction for multiple comparisons (Figure 1a). In the Inarijoki population, the only SNP that reached the Bonferroni-corrected threshold was *akap11*_Val214Met_ (Figure 1b). Specifically, the *akap11*_Val214Met_^EE^ genotype was 8.04 times (95% CI: 2.13 – 30.34) more likely to explore the nursery stream, Guoldnájohka, than the *akap11*_Val214Met_^LL^ allele (*p* < 0.001, Figure 2a). The *six6* locus did not exhibit a significant association in any of the populations.

**Figure 1:**
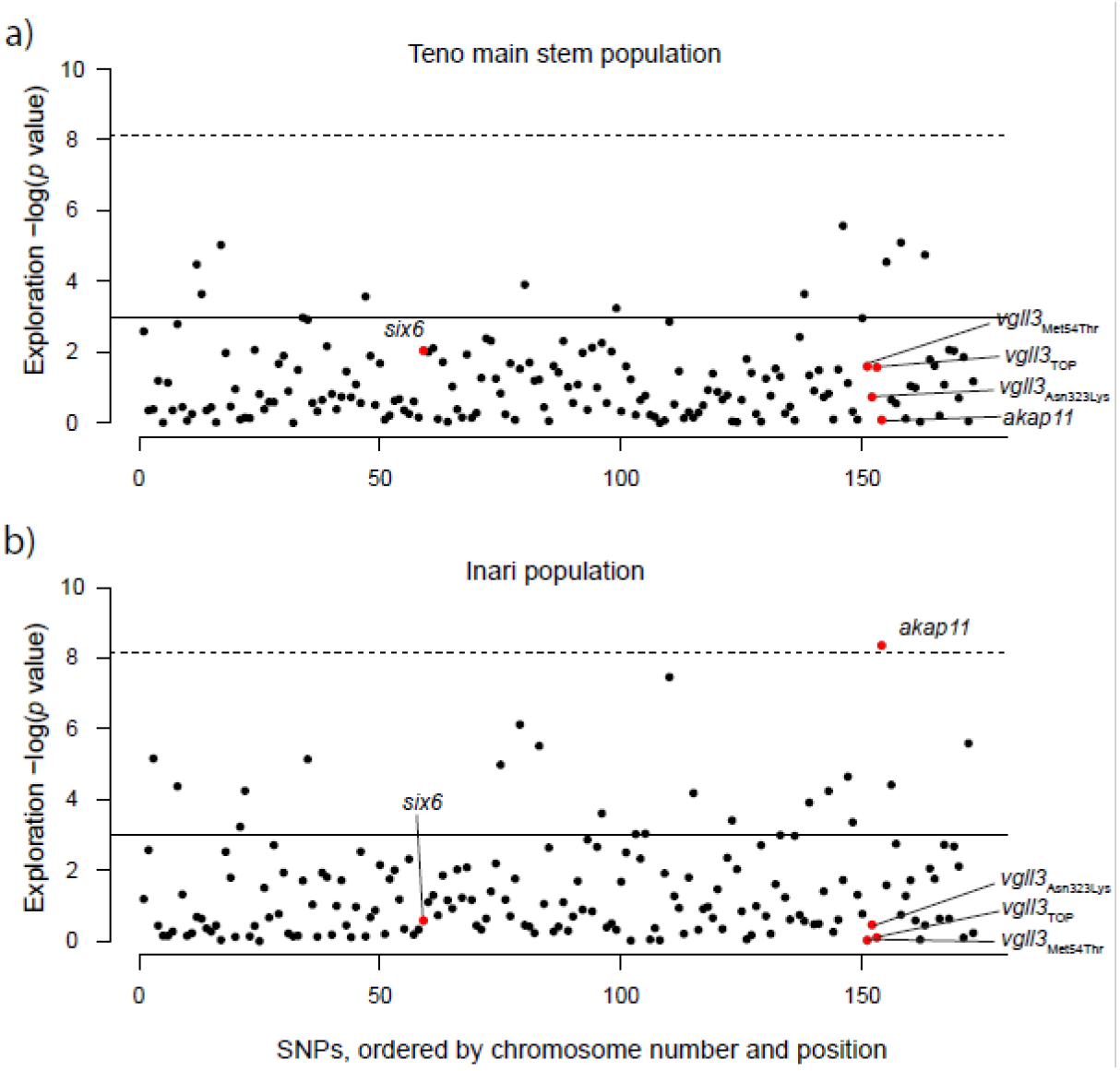
SNP - exploration association across 173 SNPs in Teno main stem (a) and Inarijoki (b) populations in Atlantic salmon. Focal SNPs are colored red. Uncorrected (*p* = 0.05) and Bonferroni corrected p-value thresholds are indicated with dashed and continuous lines.

When pairwise interactions within the *vgll3* genomic region were modeled in the Teno main stem population, none of the models were more significant than the null model that did not contain genetic effects. For the Inarijoki population, the most parsimonious model explaining the exploration included the interaction of *akap11*_Val214Met_ and *vglll3*_TOP_, which was superior to the null model (ΔAIC = 13.07), but only marginally to the model that only included *akap11*_Val214Met_ (ΔAIC = 1.12, *p* = 0.077). This model was also not significantly better than the interaction between *akap11*_Val214Met_ and two other SNPs in the *vgll3* genomic region (ΔAIC < 2, Supplementary Figure 3). This marginal interaction effect was driven by a decrease in exploration behavior associated with the genotype combination of *akap11*_Val214MetEL_ and *vgll3*_TOPEL_ (Figure 2b, Supplementary Figure 4). As such, the model predicts that the odds of exploration for individuals with *akap11*_Val214MetEL_ and *vgll3*_TOPEL_ was 3.69 (95% CI 1.63-8.36, *p*<0.001) and 3.04 (95% CI 0.80-11.47, *p*=0.14) times less than for *akap11*_Val214MetEE_ and *vgll3*_TOPEL_ and *akap11*_Val214MetEL_ and *vgll3*_TOPEL_, respectively (Figure 2b). The fit of the model with only *akap11* genotype modelled, nor the model with the *vgll3*_TOP_ and *akap11* interaction did not exhibit substantial deviation from model assumption, as assessed visually by DHARMa package’s (Hartig 2022) diagnostics plots (Supplementary Figure 5). For both rivers, exploration to nursery streams increased with age (*p*< 0.001 for both rivers), concordant with the previous reports (Erkinaro 1995).

**Figure 2:**
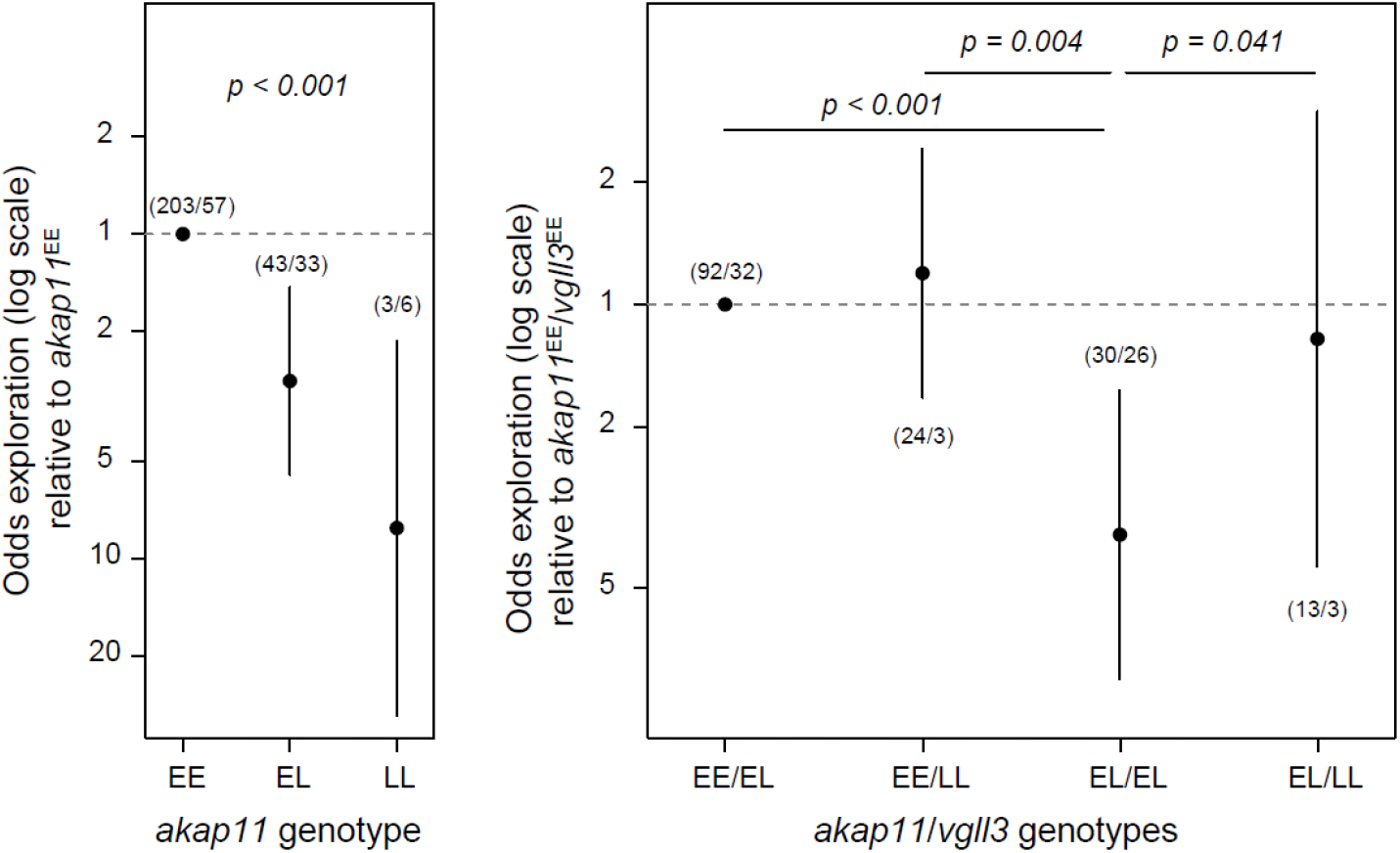
**a)** Odds of exploration in Inarijoki Atlantic salmon as a function of the *akap11*_Val214Met_ SNP, and **b)** as a function of the *akap11*_Val214Met_ and *vgll3*_TOP_ interaction. The error bars indicate 95% CI of the marginal estimates, and the first and second numbers in parentheses indicate the sample sizes in the nursery stream and the main stream, respectively. The Y axis is drawn on a logarithmic scale.

The fish sampled in Guoldnájohka contained 83 individuals that had undergone smoltification (or were in the process of becoming smolt), which is a suite of physiological, morphological, and behavioral changes in Atlantic salmon juveniles for becoming ready for seaward migration (McCormick & Saunders 1987), which may have altered their exploration behavior. However, when the smolts were excluded from the analysis, the results were mostly qualitatively similar (Supplementary Figure 6). One notable difference was that a model with the *akap11*_Val214Met_ and *vgll3*_TOP_ interaction was the best model. As in the full dataset, the fit of the interaction model the was not significantly better than the fit of the model with only additive effects of only a*kap11*_Val214Met_ (*p* = 0.12, ΔAIC = 0.19, Supplementary Figure 6). However, the change was not meaningfully different from an effect that might have resulted from a reduced sample size in a random manner. As such, when 83 parr were randomly sampled and excluded from the dataset (i.e., the same number as the smolts in the dataset) or 83 randomly selected parrs in the dataset were used as a replacement for the excluded smolts, the ΔAIC range of these models (each permuted 1000 times) was similar to the ΔAIC values obtained in the original model (Supplementary Figure 7).

## Discussion

The factors shaping the genetic architecture of ecologically important traits are fundamentally important to understand adaptive dynamics in the wild. We have demonstrated that the exploration behavior in juvenile Atlantic salmon is linked to a genomic region on chromosome 25 in one of the two populations in the Teno River. This region contains the *vgll3* and *akap11* genes and is associated with a suite of traits involved in salmon adaptation (reviewed in Mobley et al. 2021) including age at maturity (Barson et al. 2015). We have found that a missense SNP in the *akap11* gene (*akap11*_Val214Met_) was substantially linked to exploratory behavior, while SNPs in the *vgll3* gene exhibited a marginally supported epistasis with this SNP, suggesting a complex genetic architecture in the region. Furthermore, the genetic effect was only observed in one of the two populations studied, indicating the importance of environment or population-specific factors in triggering gene-dependent exploration behavior.

Our results showed that the early maturation allele in the *akap11* gene was associated with increased exploration to nursery streams in the Inarijoki population. This is consistent with the findings of Erkinaro et al. (1997) who, by analyzing freshwater growth patterns of salmon, suggested that juveniles entering nursery streams had a greater tendency to mature and return from their sea journey earlier than those that stayed in the main stem for their juvenile years. Differences in growth opportunity in Atlantic salmon juveniles, which is linked to efficient acquisition of resources, may be instrumental in exceeding the maturation threshold in subsequent years (Metcalfe 1998; Thorpe et al. 1998; Jonsson & Jonsson 2014). As such, causality between exploration and migrating to nursery streams may be suggested based on the increased likelihood of individuals in the streams to reach the maturation threshold (e.g., Thorpe et al. 1998) upon increased growth opportunities. However, the association was predominantly driven by genetic variation in the *akap11*_Val214Met_ SNP, which is about 70 kb downstream of the *vgll3* gene and exhibits a weaker association with sea age at maturity (Barson et al. 2015). The *akap11*_Val214Met_ SNP was also previously ruled out as the genetic basis for early male maturation in Atlantic salmon in controlled settings (Sinclair-Waters et al. 2022). Combined, while exploration behavior may facilitate earlier maturation in a context-dependent manner, our results do not provide evidence to support a direct genetic causation between the two, but that a small region explains both maturation and exploration behaviors, and that the two traits are genetically correlated by linkage.

A multitude of phenotypic traits may be linked within a genomic region via pleiotropy or linkage disequilibrium (that is, spurious pleiotropy, Solovieff et al. 2013). This makes it challenging to identify the phenotypic traits under selection, e.g., by using genomic scans (Paaby & Rockman 2013; Solovieff et al. 2013). Our results indicate that the *vgll3* genomic region contains multiple independent SNPs linked to adaptive traits, ruling out pleiotropy as the basis of multi-trait association. The presence of multiple adaptive traits within a tightly linked genomic region raises the question of whether the genetic covariation between these loci harbors coadapted trait combinations. Linked architectures may facilitate adaptation by enabling co-inheritance of adapted gene complexes (Mullon, Keller & Lehmann 2017; Oomen, Kuparinen & Hutchings 2020). For example, genomic inversions have been shown to be important in maintaining coadapted trait combinations, including migratory strategies and dispersal (Mullon, Keller & Lehmann 2017; Wellenreuther & Bernatchez 2018; Rubenstein et al. 2019). Similarly, genetic variation in migratory ecotypes in rainbow trout (*Oncorhynchus mykiss*, Pearse et al. 2014) and Atlantic cod (*Gadus morhua*) have been shown to be associated with genetic markers in a large inverted region (Kirubakaran et al., 2016). A tight linkage between adaptive traits might also be selected for when locally adapted populations may have varying local fitness optima (for these traits) and exhibit high gene flow between them (Yeaman & Whitlock 2011). In this case, a tight linkage between locally adapted traits would help in purging maladaptive genetic variants with a relatively reduced genetic load.

In a practical sense, our results also highlight the insufficiency of drawing conclusions when only one SNP is used as a tag marker to represent the genetic makeup of tightly linked genomic regions. In common garden cross-breeding designs, one SNP marker may be utilized to tag the whole region when exploring the trait-genotype association of the region (e.g., Prokkola et al., 2022; Bangura et al., 2024). While such designs provide substantial statistical and cost reduction benefits, they would not be able to discern associations across SNPs in close proximity, and their interactions. For example, using a common garden setup, Bangura et al., (2024), suggested that *vgll3* is likely to play a role in exploration behaviors. However, in the cross-breeding design, parents were chosen to be in complete linkage regarding the genetic variation in *akap11* and *vgll3* SNPs, thereby deduction of SNP effects independently is not possible (see also, Bangura et al., 2022). In fact, when Sinclair-Waters et al., (2022) refined the association of SNPs in the vgll3 genomic region, in relation to their effect on male parr maturation, they found an opposite effect of SNPs in the *vgll3* and *akap11* gene, whereby the early maturation allele in the *akap11* gene was indeed marginally associated with reduced maturation, further signifying the importance of refining genetic associations when multiple candidate SNPs may underlie the phenotypic variation.

Our study provides a further outlook on the dynamics of behavioral syndromes and correlated behaviors (Höjesjö, Johnsson & Bohlin 2004; Garamszegi, Markó & Herczeg 2013; Mathot, Dingemanse & Nakagawa 2019; Axling et al., 2023). The classic paradigm suggests that dispersal is negatively correlated with dominance and aggression, whereby dominant individuals monopolize favorable habitat patches and force subordinate individuals to displace in search of new habitats (Stamps 1994; Stamps 1999). However, in Atlantic salmon, migration to nursery streams might be carried out by individuals holding higher positions in the dominance hierarchy (Erkinaro & Niemelä 1995). Using seasonal growth in scale depositions, Erkinaro and Niemelä (1995) estimated growth in previous seasons, a trait correlated with dominance and aggression in early life stages (Cutts et al., 2005), and showed that faster growing individuals are more likely to migrate to the small nursery streams. A similar tendency in individual growth patterns has also been identified in individuals shifting their juvenile rearing habitat to lacustrine areas of the catchment (Erkinaro et al., 1998). A niche shift, i.e., by migrating to nursery streams, may be hypothesized to be a mechanism to overcome potential resource mismatch in the original habitat. Therefore, we would predict that the EE allele in the *vgll3* genomic region is likely associated with behavioral traits correlated with exploration in salmon, such as dominance.

On the other hand, common garden experiments under controlled conditions do not support our prediction. For example, by quantifying the exploration of new environments in a confined space (an aquarium with a surface area of 25 × 30 cm), Bangura et al., (2024) found that the *vgll3* L allele was associated with greater exploration in a novel environment. Similarly, using a similar experimental setup, Bangura et al., (2022) also found increased aggression associated with the *vgll3* L allele. These results suggest that there is a genetic covariation between aggression and novel environmental exploration, but with an opposite direction of allelic effect to our study. (Note that in the crosses used by Bangura et al., (2022) and Bangura et al., (2024) *akap11* and *vgll3* were in full linkage, so conclusions related to the genetic variation in the *vgll3* SNPs are equally valid for the *akap11* SNP.) These contrasting results can reflect contextual differences between natural and common garden settings. As such, a meta-analysis has shown that these two personality traits (aggression and exploration) only correlate when there is a contextual overlap that underlies the similarity of trait expression (Garamszegi, Markó & Herczeg 2013). Therefore, the contrasting conclusions in these two settings (natural vs. laboratory) may stem from different functional physiological processes. Alternatively, strong environment (common garden vs. natural setting) by genotype, or population by genotype interactions may explain the contrast. For the latter, for example, the parental fish used in Bangura et al., (2022) and Bangura et al., (2024) are obtained from a hatchery broodstock of salmon managed by the Natural Resources Institute Finland (LUKE), initially originated from a Baltic Sea population (River Neva). Therefore, differences in the genetic background of fish may also explain the discrepancies. In fact, the genetic association between SNPs in the *vgll3* genomic region and exploration was significant only in the Inarijoki population, while no detectable effect was found in the Teno main stem population, further highlighting a potential role of population-specific differences in the genetic basis of trait variation. Our study illustrates the complex dynamics of the correlated behaviors of Atlantic salmon and their genetic background.

Finally, we found a marginal support for epistasis between *akap11* and the *vgll3* gene, in which the fit of the interaction model was better than the fit model with only *akap11* SNP, but this did not reach to a significance at alpha value = 0.05. The statistical power was also insufficient to determine which of the SNPs in the *vgll3* gene are the most likely candidates. However, the suggestive signals of epistasis may be an indication of a complex genetic architecture within the genomic region. Epistasis contributes to the trait variance and helps maintain additive genetic variation upon selection (Cheverud & Routman 1995). Furthermore, the trait complex between exploration and age at maturity may contribute to the adaptation dynamics. For example, optimal trait combinations (of traits co-inherited within a haplotype block) may be locally variable among populations and may improve overall fitness compared to what would be predicted by only additive gene action. Such an epistatic interaction would also facilitate purging of maladapted genetic combinations (Yeaman & Whitlock 2011).

We used historical field data that were not originally intended for a genetic study such as this. Therefore, the design has some drawbacks. For example, the hatching year, which represents the year that the offspring originate from, was not modelled for the Teno main stem and Baððá stream due to the lack of temporal overlap between sites. Therefore, we cannot rule out the possibility that the (lack of) difference in genetic response is due to temporal changes in allele frequencies between cohorts, even though the time span between sampling periods is only a few years. However, the hatch year was accounted for in the model for the Inarijoki population, by which we controlled for temporal genetic changes between cohort years. Second, genotype calling failed for a substantial portion of individuals due to the suboptimal amount of tissue remaining in the storage envelopes. In addition, we used a targeted genotyping approach by scoring only the most likely candidate SNPs in the genomic region. An alternative approach to sequence the whole genomic region might provide finer resolution in trait association and help to better narrow down the causal region.

In this study, two life history genomic regions were the primary focus (*vgll3* and six6), and other SNPs genotyped was used to account for potential inflation in the statistical tests. Therefore, by design, we did not perform large scale scans to explore the signals of association at genomic scale, which may provide insights to the molecular mechanisms underlying the trait variation. Genome wide association analyses have identified a number small effect loci linked to age-at-maturity (Ayllon et al., 2015; Gutierrez et al., 2015; Sinclair-Waters et al., 2020), but such studies for traits such as movement or exploration are mostly missing in non-model organisms, e.g., Atlantic salmon. Theory of omnigenic genetic architecture predicts that major regulatory proteins may drive covariation in multiple traits via highly pleiotropic function (Boyle et al., 2017; Liu et al., 2019). In humans, genome-wide meta-analyses have been used to genetically link personality traits to larger psychological disorders (Lo et al., 2017; Gupta et al., 2024). Such approaches as in human studies may help to better understand to linking exploration and life-history at the molecular level and potential regulatory networks shared by these two traits (Liu et al., 2019; Boyle et al., 2017; Mathieson 2021), once the genome-wide association are available for the trait of interests (i.e., exploration).

Our system included only two populations from a northern River which are generally low-nutrient, low productivity areas with depauperate fish species communities, and whether the observed genetic effects are parallel across the species range is unclear. The association between age-at-maturity and the *vgll3* genomic region is sex and region specific for the North American populations (Kusche et al., 2017; Kess et al., 2024), and may have an environmental component in some European populations (Bersnier et al., 2019; Ayllon et al., 2019). Exploration behavior may alter life-history variation context dependently, e.g., via means of enabling improved feeding opportunities (Wright et al., 2007; Chapman et al., 2012). Therefore, it is intriguing to ask whether such context dependent association between the *vgll3* genomic region and age-at-maturity may have linked to the *akap11* gene.

Movement patterns in the Salmonidae family are mostly studied in the context of the phenology of migration, which is central to their life histories, has a heritable component, and has strong fitness consequences (Niemelä et al., 2006; Stewart, Middlemas & Youngson 2006). Many species exhibit a strong dichotomy of the trait within and among populations, as evidenced by the presence of both resident and migratory forms in, e.g., brown trout, rainbow/steelhead trout, and Arctic char (Jonsson & Jonsson 1993). Movement within the freshwater habitat at the juvenile stage and its association with migration at later stages are much less explored. Our study demonstrates a complex genetic interaction between freshwater exploration and maturity age. The linked inheritance of these two functional traits (exploration and maturation) may help to maintain coadapted trait complexes, and may be instrumental in local adaptation processes, fitness, and the long-term survival of populations.

## Acknowledgement

We thank Susanna Ukonaho and Meri Lindquist from the University of Turku for help with DNA extraction, Matti Kylmäaho from Luke for scale aging, and Lars Paulin from the University of Helsinki for coordinating the genotyping-by-sequencing assays. Funding came from the Research Council of Finland (353388 and 325964 to T.A.). We thank to Craig Primmer for his communication during the making of the study.

