## Supplementary Materials for "Early life exploration behavior and life-history loci are co-localized in an adaptive genomic hotspot in Atlantic salmon"

**Supplementary Table 1:** Age and hatch year of individuals used in this study across populations (Teno, Inarijoki) and their main stems and small tributary (nursery) streams. The numbers in parenthesis are smolts.

| | | | Hatch year | | | | | |
|---|---|---|---|---|---|---|---|---|
| | | Age | 1988 | 1989 | 1990 | 1991 | 1992 | 1993 |
| Teno | main stem | 1 | 0 | 0 | 10 | 0 | 0 | 0 |
| | | 2 | 0 | 36 | 0 | 0 | 0 | 0 |
| | | 3 | 21 | 0 | 0 | 0 | 0 | 0 |
| | Baðđá | 1 | 0 | 0 | 0 | 0 | 5 | 50 |
| | | 2 | 0 | 0 | 0 | 6 | 52 | 0 |
| | | 3 | 0 | 0 | 5 | 18 (4) | 0 | 0 |
| | | 4 | 2 (2) | 9 (5) | 44 (43) | 0 | 0 | 0 |
| | | 5 | 18 (17) | 14 (14) | 0 | 0 | 0 | 0 |
| Inarijoki | main stem | 1 | 0 | 0 | 0 | 0 | 14 | 5 |
| | | 2 | 0 | 14 | 0 | 21 | 15 | 0 |
| | | 3 | 16 | 0 | 5 | 6 | 0 | 0 |
| | Guoldnájohka | 1 | 0 | 0 | 15 | 4 | 3 | 22 |
| | | 2 | 0 | 28 | 0 | 12 (1) | 34 | 0 |
| | | 3 | 4 | 0 | 8 | 7 | 0 | 0 |
| | | 4 | 0 | 61 (28) | 51 (48) | 0 | 0 | 0 |

**Supplementary Table 2**: Observed number of individuals in the Inarijoki population, main stem (0) vs the Guoldnájohka stream (1).

| *akap11* | *vgll3*$_{TOP}$ | 0 | 1 |
|---|---|---|---|
| EE | EE | 22 | 87 |
| EE | EL | 32 | 92 |
| EE | LL | 3 | 24 |
| EL | EE | | 0 |
| EL | EL | 26 | 30 |
| EL | LL | 3 | 13 |
| LL | EE | 0 | 0 |
| LL | EL | 4 | 1 |
| LL | LL | 2 | 2 |

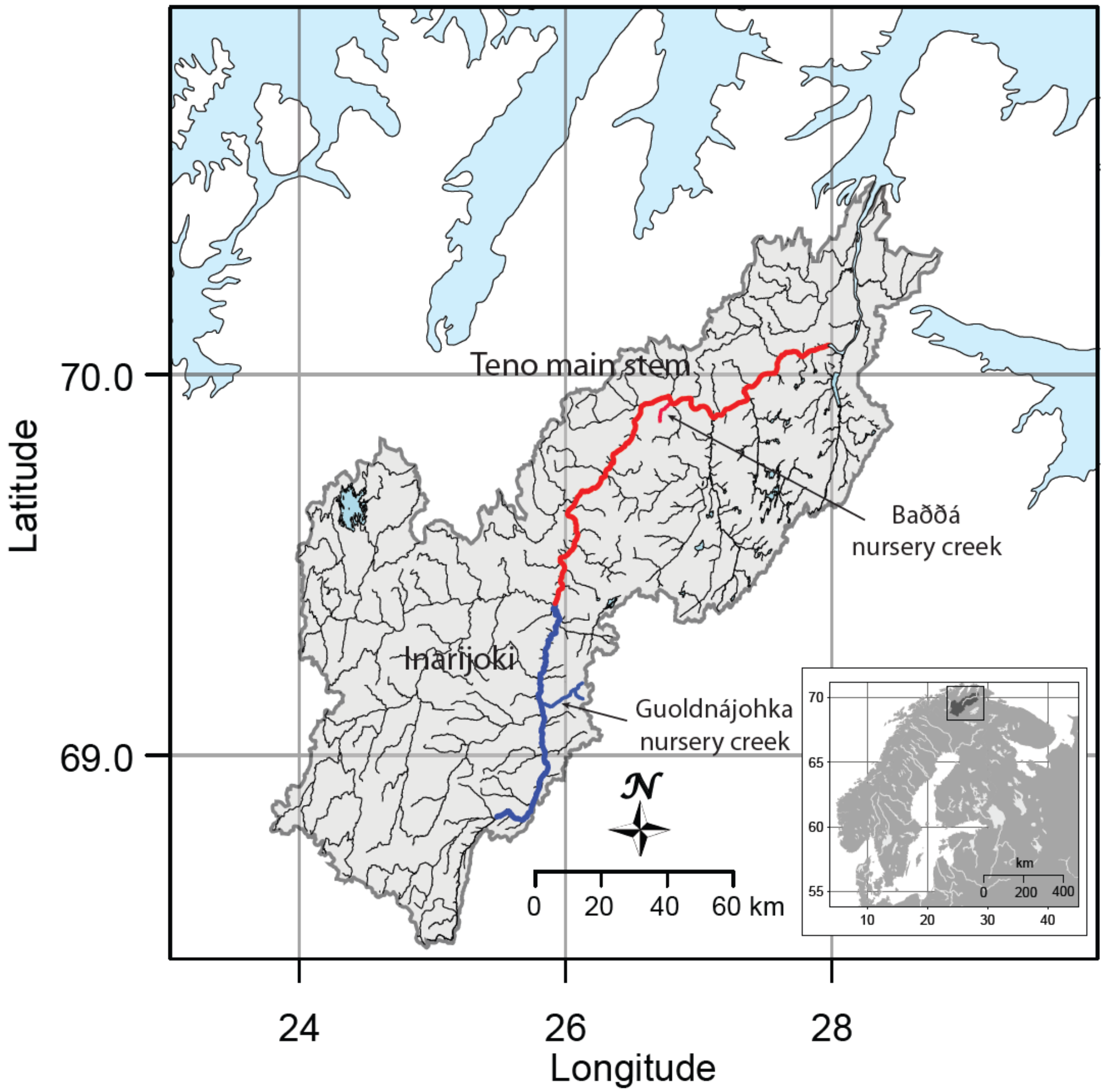


**Supplementary Figure 1:** The study system and the Teno River Basin. River sections with blue and red colors indicate Teno main stem and Inarijoki, respectively, and respective nursery streams, Baðđá and Guoldnájohka. Note that the sampling in both spawning rivers were carried out in close proximity (up to 100 meters) to the outlets of two nursery streams.

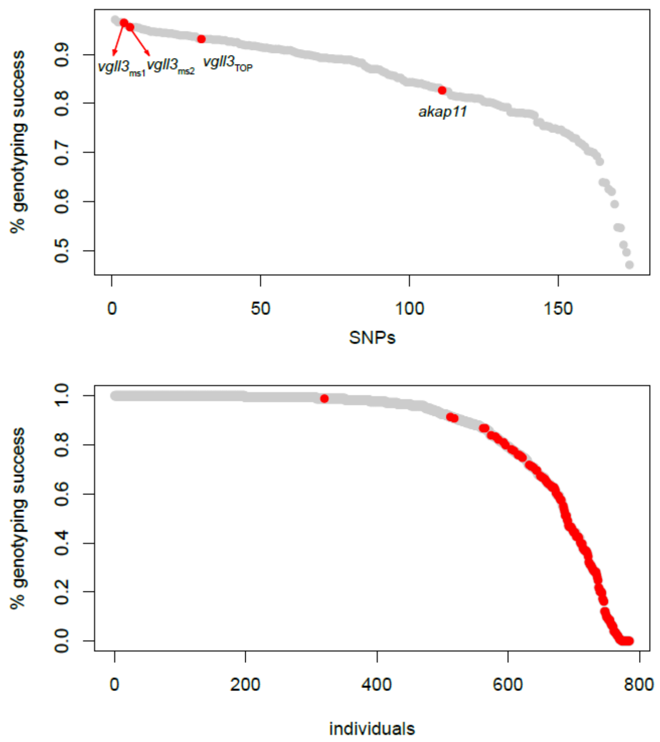


**Supplementary Figure 2:** Genotyping success per individuals (a) and SNPs (b). Focal SNPs are marked in red in (a). Excluded individuals were marked in red in (b).

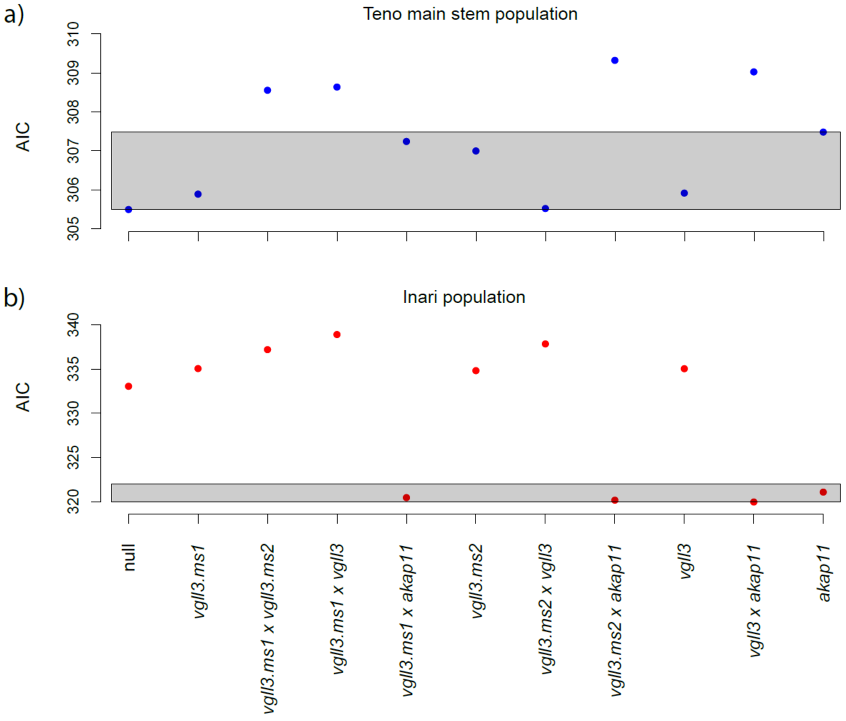

**Supplementary Figure 3:** Parsimony of different model structures as evaluated by Akaike information criteria (AIC). Models in the gray shaded area indicates similarly parsimonious model (i.e., models within 2 AIC units to the best model.)

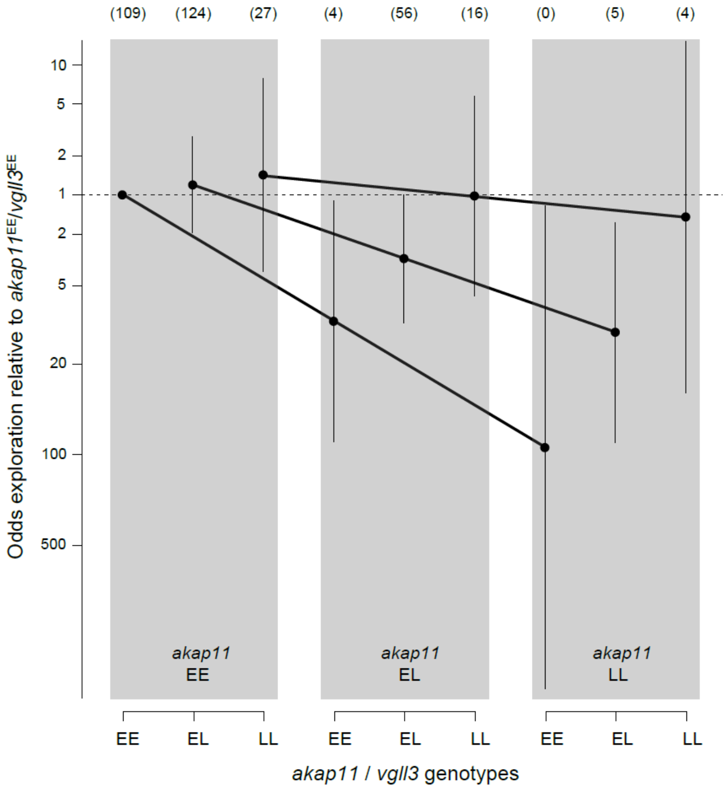


**Supplementary Figure 4:** Predicted odds of exploration in Inarijoki Atlantic salmon as a function of *akap11* and *vgll3* interaction with all genotype combinations are visualized. Error bars indicate 95% CI of the marginal estimates and numbers in parenthesis indicates sample size. Y axis is drawn in log scale.

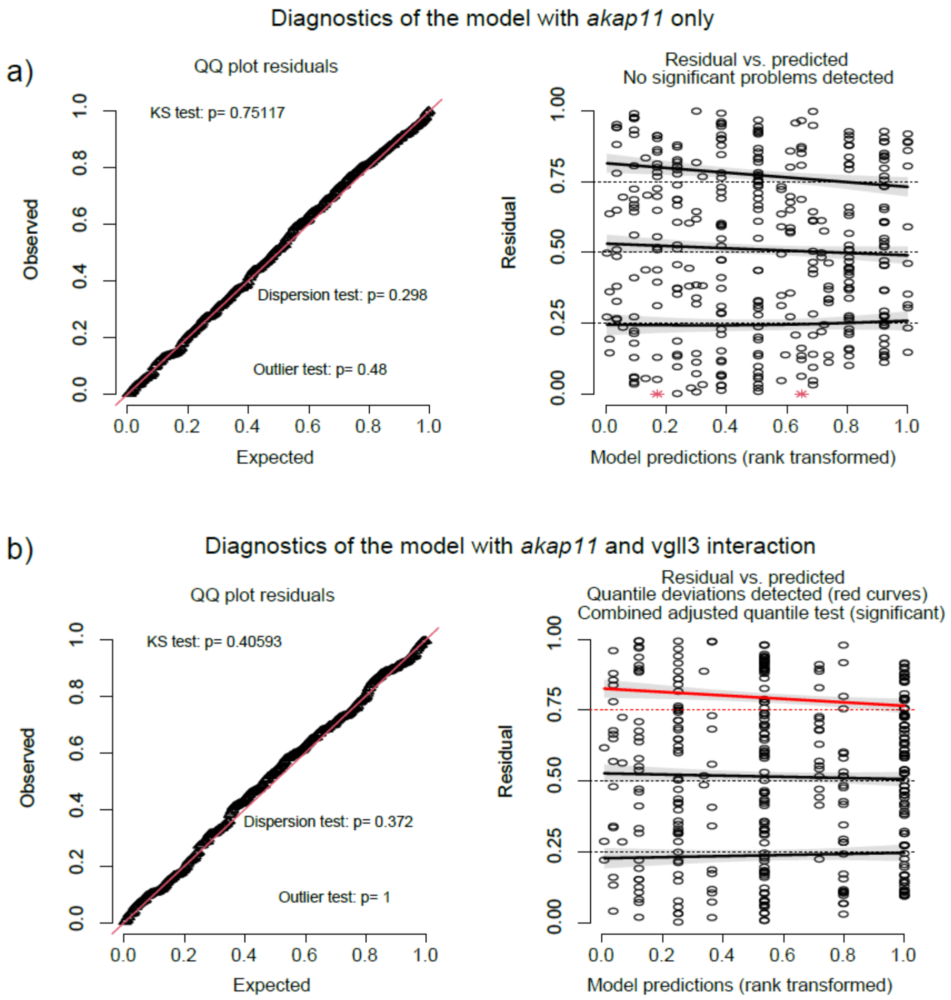


**Supplementary Figure 5:** Diagnostic plots of the models that has only *akap11* as the genotype factor (a), and that modelled *akap11* and *vgll3* interactions.

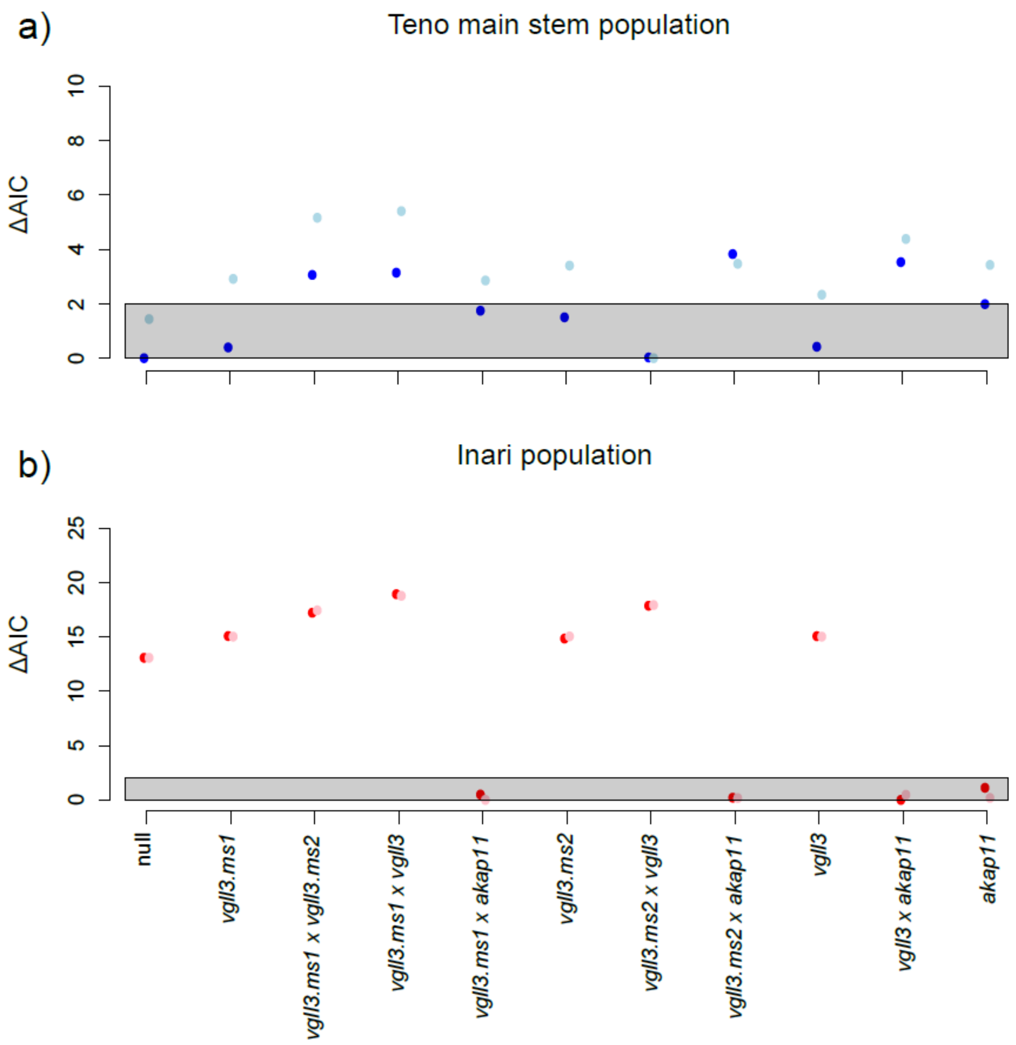


**Supplementary Figure 6:** Comparing the parsimony of model structures between modelled with and without smolts included in the dataset (dark *vs.* light symbols, respectively), as evaluated by Akaike information criteria (AIC). Models in the gray shaded area indicates similarly parsimonious model (i.e., models within 2 AIC units to the best model.)

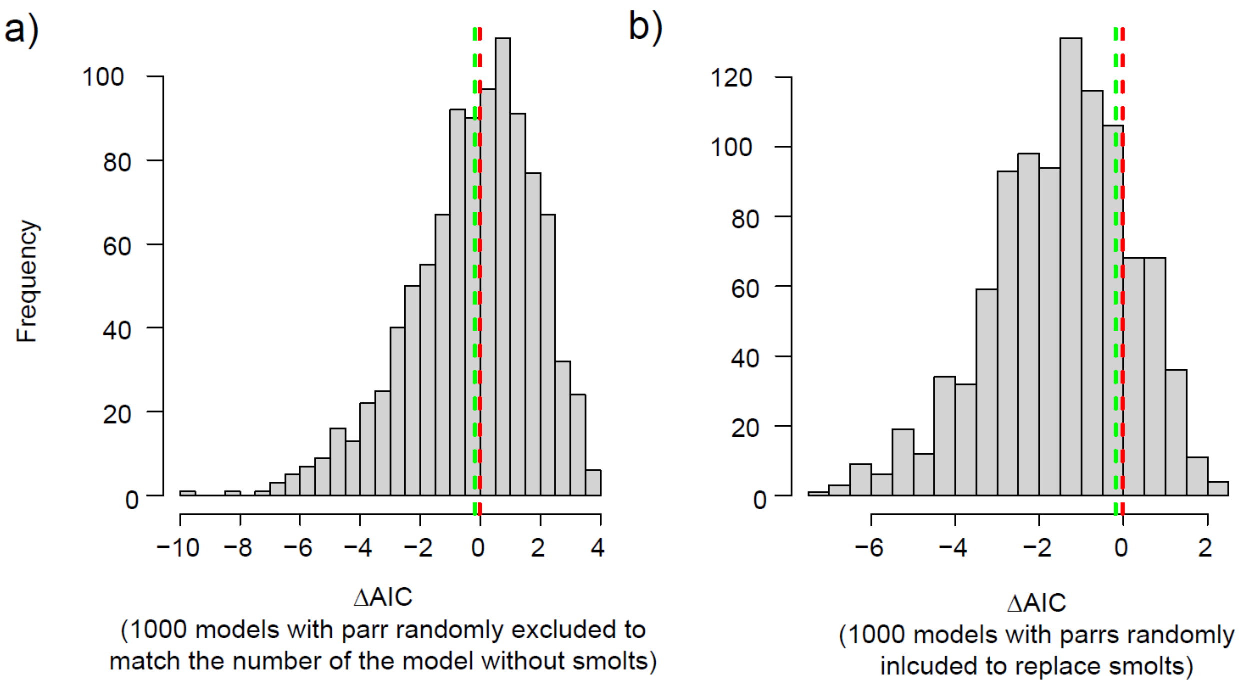


**Supplementary Figure 7:** Evaluating 1000 permutated models to evaluate the account of individuals that had undergone parr-smolt transformation (smolts) in the dataset when evaluating the model parsimony (ΔAIC) between *akap11* additive model vs *vgll3*$_{TOP}$ and *akap11* interaction model. In (a), same number of parrs as the number of smolts in the data were excluded from the dataset, and parsimony between two model structures were compared, i.e., between the real model that excludes smolts (red dashed lines) and 1000 permuted models. In (b), same number of parrs as the number of smolts in the data were included to the dataset, and parsimony between two model structures were compared, i.e., between the real model that still include smolts (red dashed lines) and 1000 permuted models.
